# A Self-Limiting Circuit Regulates Mammary Cap Cell Plasticity Through TGF-beta Signaling

**DOI:** 10.1101/2021.08.04.455067

**Authors:** Armelle Le Guelte, Ian G. Macara

## Abstract

The organization and maintenance of complex tissues requires emergent properties driven by self-organizing and self-limiting cell-cell interactions. We examined these interactions in the murine mammary gland. Luminal and myoepithelial subpopulations of the postnatal mammary gland arise from unipotent progenitors, but the destiny of cap cells, which enclose terminal end buds (TEB) in pubertal mice, remains controversial. Using a transgenic strain (Tg11.5kb-GFP) that specifically marks cap cells, we found ~50% of these cells undergo divisions perpendicular to the TEB surface, suggesting they might contribute to the underlying luminal cell population. To address their stemness potential we developed a lineage tracing mouse driven from the s-SHIP (11.5 kb) promoter. Induction of tdTomato (tdTom) from this promoter in vivo demonstrated that all cap cell progeny are myoepithelial, with no conversion to luminal lineage. Organoid cultures also exhibited unipotency. However, isolated cap cells cultured as mammospheres generated mixed luminal/myoepithelial spheres. Moreover, ablation of luminal cells in vivo using diphtheria toxin triggered repopulation by progeny of tdTom+ cap cells. A signaling inhibitor screen identified the TGFβ pathway as a potential regulator of multipotency. TGFβR inhibitors or gene deletion blocked conversion to the luminal lineage, consistent with an autocrine loop in which cap cells secrete TGFβ to activate the receptor and promote luminal transdifferentiation. Ductal tree regeneration in vivo from isolated cap cells was much more efficient when they were pre-treated with inhibitor, consistent with more cells retaining cap cell potential prior to transplantation. Notably, in vitro transdifferentiation of cap cells was blocked by co-culture with luminal cells. Overall, these data reveal a self-limiting cell circuit through which mammary luminal cells suppress cap cell conversion to the luminal lineage.

## INTRODUCTION

The murine mammary gland provides a powerful model system for the investigation of self-organization of complex tissues, and how collective decisions determine cell fate. A ductal tree emanating from the nipple is elaborated during puberty and is composed of polarized luminal epithelial cells expressing E-cadherin (E-cad) and Keratin 8 (K8), enclosed by a layer of Keratin 14 (K14) positive basal myoepithelial cells (Silberstein, 2001). Ductal outgrowth occurs from terminal end buds (TEBs) that invade the mammary fat pad and which consist of luminal body cells surrounded by a single layer of basal-like cap cells (Wiseman et al., 2003). Both myoepithelial and cap cells can efficiently regenerate complete, functional ductal trees when transplanted into the mammary fat pads of recipient mice (Bai and Rohrschneider, 2010; Shackleton et al., 2006; Stingl et al., 2006); and isolated myoepithelial cells in culture can begin to express luminal markers, demonstrating a multipotent potential l(Van Keymeulen et al., 2011). But although lineage-tracing studies have shown that embryonic mammary cells are multipotent, the postnatal luminal and basal populations arise exclusively from unipotent progenitors (Van Keymeulen *et al.*, 2011; Wuidart et al., 2016; Wuidart et al., 2018). Why would basal cells retain a capacity for multipotency that is not utilized postnatally? One possibility is that these cells can switch lineage to engage in tissue repair after wounding. Indeed, we have shown recently that DNA damage triggers myoepithelial lineage switching to generate hyper-proliferative luminal cells (Seldin and Macara, 2020); and the ablation of luminal cells by diphtheria toxin activates multipotency in the myoepithelial population, potentially through multiple signaling pathways (Centonze et al., 2020). However, whether cap cells are unipotent and possess the same capacity for lineage plasticity remains controversial. Ballard et al (Ballard et al., 2015) have reported that cap cells can undergo asymmetric cell divisions, but lineage tracing with an inducible reporter driven from the smooth muscle actin promoter or from the p63 promoter (which are both expressed in cap cells as well as in myoepithelial cells) did not detect any contribution to the luminal cell population (Abdul-Manan et al., 1999; Prater et al., 2014).

To rigorously address this issue, we generated a new lineage tracing mouse using the s-SHIP promoter to drive a CreER fusion protein, which on treatment with tamoxifen triggers tdTom expression from the Rosa26 locus by removal of an upstream transcription termination sequence. The s-SHIP 11.5 kb promoter is active only in cap cells within the pubertal mammary gland, and in alveoli of pregnant mice (Bai and Rohrschneider, 2010). We found that tdTomato (tdTom+) cells exclusively generate myoepithelial progeny both in puberty and pregnancy. However, expression of diphtheria toxin receptor (DTR) from a K8 promoter and injection of toxin into the mammary glands to ablate the luminal population, induces a switch by cap cells to the luminal lineage. Similarly, isolated cap cells grown in 2D culture gave rise to cells expressing luminal markers. Strikingly, co-culture with luminal cells blocked this lineage switch. Moreover, we discovered that TGFβ signaling promotes transdifferentiation. We propose that TGFβ acts as an autocrine signal, suppressed by adjacent luminal cells, to promote luminal conversion. This self-limiting cell circuit explains why in the unperturbed state the cap cells behave as unipotent myoepithelial progenitors but can generate luminal cells when isolated or when neighboring luminal populations are damaged, thereby maintaining tissue homeostasis.

## RESULTS

### Cap cells are unipotent myoepithelial lineage progenitors

Based on previous data suggesting that cap cells can undergo asymmetric divisions (Ballard *et al.*, 2015), we first determined whether they divide only parallel to the surface of the TEB or also show perpendicular divisions. The s-SHIP 11.5kb-GFP transgenic mouse was used as this strain specifically marks cap cells with GFP (Bai and Rohrschneider, 2010). We found that these cells show a random distribution with about half the GFP+ cells dividing parallel and half dividing roughly perpendicular to the TEB surface (Fig 1A, B). This result is consistent with observations of occasional GFP+ cells within the TEB body (Fig 1A) (Bai and Rohrschneider, 2010). Previous studies have demonstrated that, rather than converting to luminal cells, however, these internalized cap cells undergo apoptosis (Sreekumar et al., 2017). The reason for this process remains unknown.

**Figure 1.**
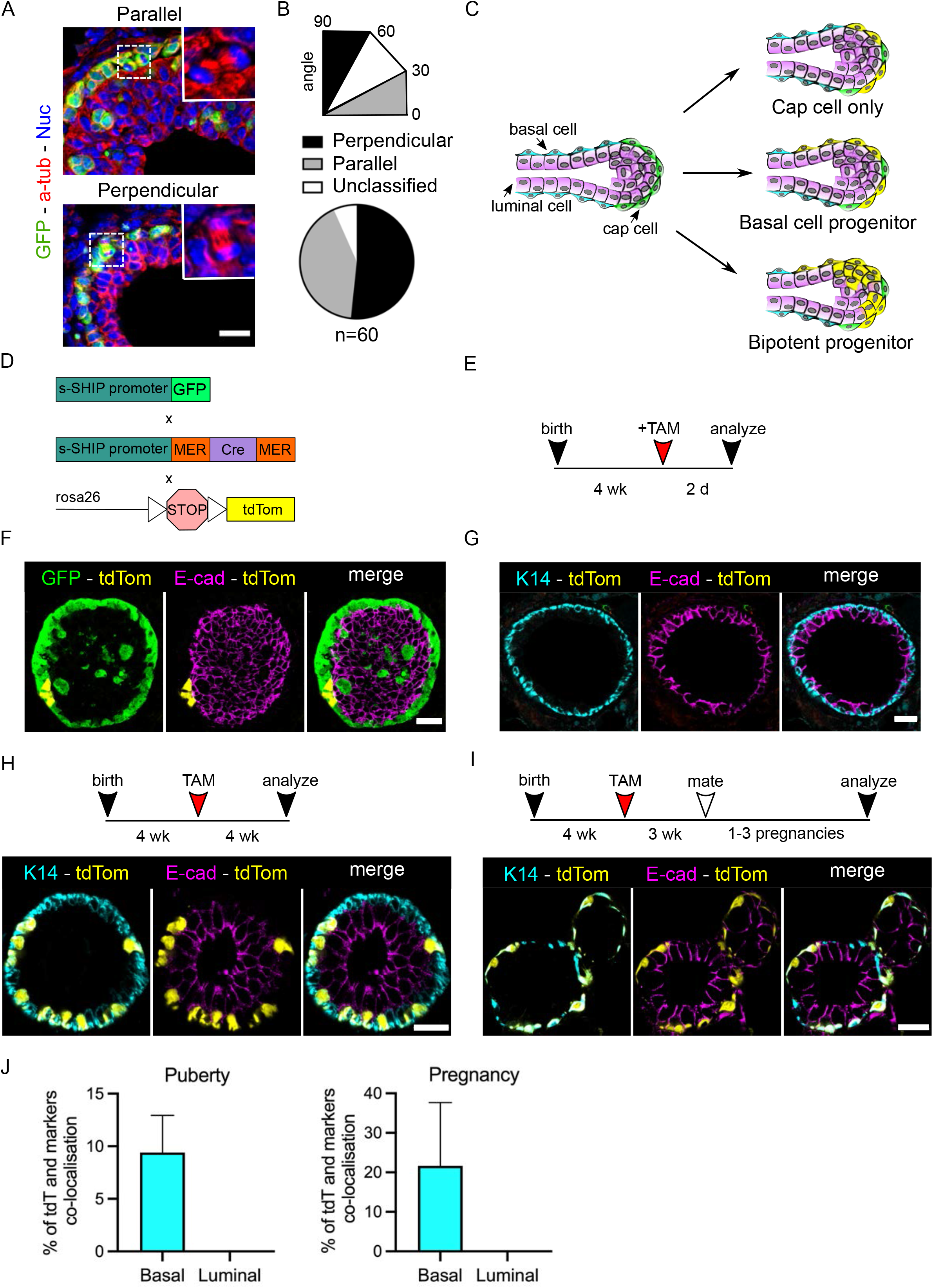
Cap cells contribute to the basal lineage. A. Representative image of parallel and perpendicular dividing cap cells in 6 week-old mammary tissue sections obtained by immunostaining of Hoechst, α-tubulin and GFP. Scale bar 20 μm. B. Quantification of cap cell division angles relative to the basement membrane. n=60 C. Graphic representation of different hypothetical outcomes of cap cell division in the TEB. D. Schematic summarizing the genetic strategy used to lineage trace cap cells and their progeny. E. Protocol to study cap cells fate at different stages of development. F-I. Immunostaining of GFP (green), K14 (cyan), E-cad (magenta) and tdTom (red) in mammary glands during puberty (2 days or 4-week post TAM injection) and at 14.5 days of pregnancy. n=3 independent experiment each. F-I. Scale bar 20 μm. J. Quantification of the number of tdTom^+^ cells for each epithelial lineage at puberty and pregnancy.

We next asked if cap cells simply self-renew but retain their identity, or function as myoepithelial cell progenitors, or instead act as bipotent progenitors that can give rise to luminal as well as basal cells (Fig 1C). To rigorously distinguish these three hypotheses, we created a new lineage-tracing mouse using the s-SHIP 11.5kb promoter to drive a CreER fusion protein, with a lox-STOP-lox cassette upstream of tdTom driven from a CAG promoter at the Rosa26 locus (Fig 1D). For some experiments this strain was crossed to the Tg11.5kb-GFP mouse to mark all the cap cells green. The tdTom tracer was activated in cap cells by treatment of the mice at 4 weeks of age with Tamoxifen to activate the CreER fusion protein. Conversion was highly specific to the cap cells (Fig 1E–G; Supp Fig 1A, B). No luminal cells or mature myoepithelial cells (K14+) were marked by tdTom when glands were examined 2 days after activation of the tracer (Fig 1F-G). However, when the analysis was performed 4 weeks post treatment with Tamoxifen, myoepithelial (K14+) cells along the ducts were labeled with tdTom, although luminal cells were still not labeled (Fig 1H; Supp Fig 1C). Notably, two distinct founders gave similar results in terms of specificity and efficiency (Supp Fig 1A-C).

As the s-SHIP promoter is also active in alveolar basal cells we performed lineage tracing on lactating mice after inducing tdTom expression prior to mating. Again, only K14+ basal cells were labeled with tdTom (Fig 1I, J, Supp fig 2). The CreER-mediated recombination was fairly inefficient, with about 9% of cells expressing the label during puberty and 20% during pregnancy (Fig 1J). We conclude, therefore, that cap cells function not as stem cells but as self-renewing unipotent progenitors of the myoepithelial lineage.

Does the cap cell population contribute significantly to ductal elongation? Mature myoepithelial cells can continue to proliferate in the ducts, suggesting that cap cells might not be essential (Giraddi et al., 2015). To test their functional significance in mammary gland development, we crossed the s-SHIP promoter-CreER line to a lox-STOP-lox DTA mouse (Supp Fig 3A). Addition of tamoxifen should induce diphtheria toxin specifically in cap cells, killing them but not pre-existing myoepithelial cells. As shown in Supplementary Fig 3 B-D, outgrowth of ducts into the mammary fat pat was significantly reduced by tamoxifen treatment of the CreER;DTA mice, as compared to tamoxifen treatment of DTA mice lacking cap-cell specific CreER expression. Note that tamoxifen treatment alone reduces outgrowth, which is a known side effect of this estrogen receptor antagonist (Hovey et al., 2005) (Supp Fig 3C, D). A 4-fold decrease of number of cap cells was observed by FACS after cap cell ablation (Supp Fig 3E, F). We conclude that cap cells contribute to ductal outgrowth, presumably by generating new myoepithelial cells, though they likely also contribute by expression of axonal guidance proteins and secretion of proteases that degrade the extracellular matrix ahead of the terminal end buds (Morris et al., 2006; Wiseman *et al.*, 2003).

### Cap cells can generate luminal-like cells in vitro

Van Kaymeulen et al demonstrated that myoepithelial cells grown in culture could begin to express luminal markers (Van Keymeulen *et al.*, 2011). Therefore, we asked if cap cells can also convert to the luminal lineage when isolated from the mammary niche. Initially, we generated organoids, using our lineage-tracing mouse, and found that they recapitulated the in vivo phenotype, with tdTom expressed in the myoepithelial cells (marked with K5 or p63) but with no labeling of luminal cells (identified by E-cad staining) (Fig 2A, B).

**Figure 2.**
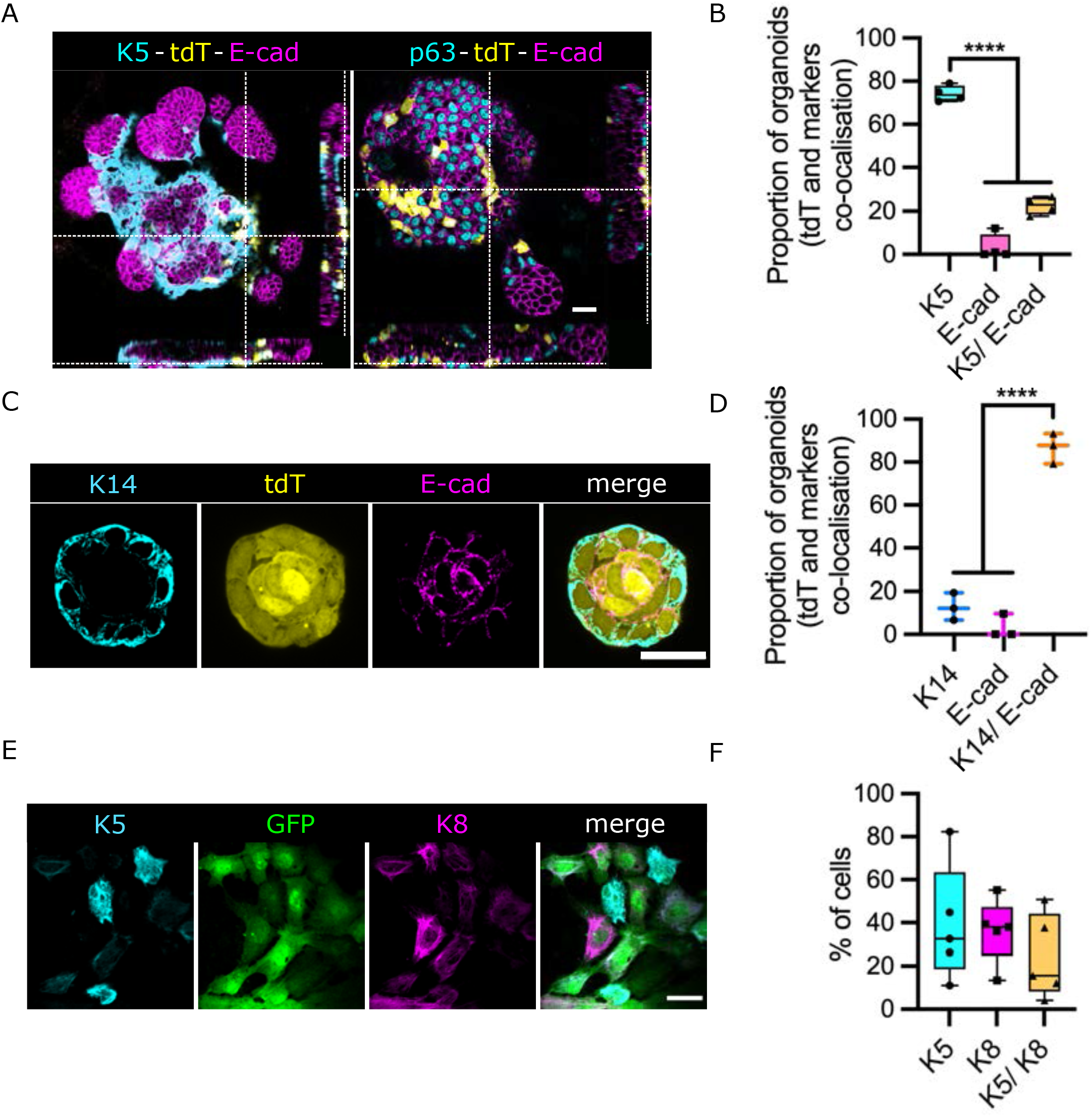
Disruption of the micro-environment promotes cap cell transdifferentiation. A. 3D and lateral images of organoids grown in 3D Matrigel culture from s-SHIP-GFP/s-SHIP-Cre-tdTom mice and immunostained for K5, E-cad and tdTom. Organoids were treated with 4-OH Tam and grown for 7 days prior to staining. B. Quantification of cell types in organoids labeled with the tdTom cap cell lineage tracer. n=4 independent experiments C. Mammospheres isolated from single cells, treated with 4-OH-Tam and immunostained for K14 (cyan), E-cad (Magenta) and tdTom (yellow). D. Quantification of tdTom^+^ mammosphere cell lineage composition. n=3 independent experiments. E. 2D culture of FACS isolated s-SHIP-GFP cells. Images were taken after 1 week and cells were stained for K5 (cyan), GFP (green) and E-cad (magenta). F. Proportions of luminal and basal cells were quantified at day 4 (note all values were zero at t = 0 days). n=3 independent experiments. A., C. and E. Scale bar = 20 μm. p-Values were determined by Sidak’s multiple comparison test. (*p< 0.05, **p< 0.01, ***p< 0.001, ****p< 0.0001).

Strikingly, however, when single cells from the lineage tracing mouse were isolated by FACS (Supp Fig 3F) and grown in Matrigel as mammospheres, tdTom+ colonies exhibited labeling of both myoepithelial (K14+) and luminal (E-cad+) cells (Fig 2C, D). Moreover, purified GFP+ cap cells grown in 2D cultures also gave rise to K8+ luminal cells as well as K5+ basal cells (Fig 2E, F). These results suggest that organoid cultures maintain the mammary gland micro-environment to effectively suppress conversion of the cap cells to the luminal lineage, but that in mammospheres grown from single cells and in 2D cultures this restrictive environment is lost.

### Ablation of luminal cells in vivo permits cap cell multipotency

To test if this conversion to luminal cells is exclusive to in vitro culture or can occur in the in vivo setting, we asked if the ablation of luminal cells in mammary ducts of pubertal mice would, by destroying the local micro-environment, permit cap cells to transdifferentiate into new luminal cells, to repair the damage. With this goal, we crossed our lineage-tracing mouse to a K8-GFP-DTR mouse, which expresses DTR plus a GFP marker specifically in luminal cells (Fig 3A, B). The lineage marker tdTom was induced in these mice by tamoxifen treatment at 4 weeks of age; DTA was then injected into the mammary fat pads at 5 weeks, and the mammary glands were analyzed at 6 weeks (Fig 3 B, C). GFP was expressed in luminal cells throughout the ducts in the absence of DTA; however, many GFP+ cells were ablated by injection of the toxin (Fig 3C, Supp Fig S4). An apoptotic marker, cleaved Caspase 3, was detectable only in the luminal cells of mice treated with DTA; and strikingly, tdTom+/Ecad+ dual-positive cells were also detectable in these mice, demonstrating that luminal cell damage promotes conversion of cap cells to the luminal lineage (Fig 3C, Supp Fig S4).

**Figure 3.**
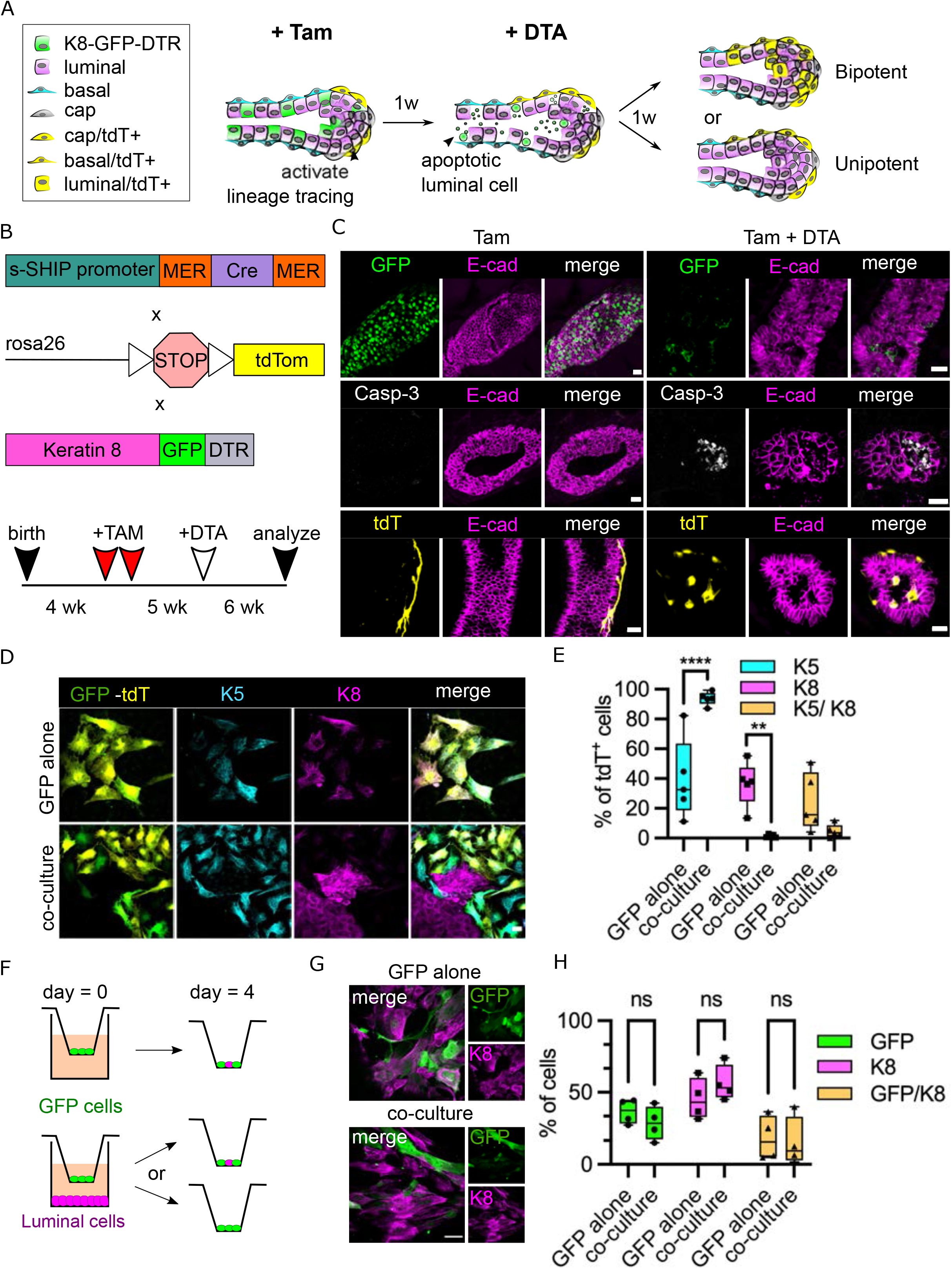
Ablation of luminal cells in vivo permits cap cell multipotency and a short-range signal from luminal cells blocks cap cell multipotency. A. Graphic representation of approach to test effect of luminal cell ablation on cap cell fate. B. Protocol of the strategy used to track cap cells progenies after luminal cell ablation. C. Confocal images of ductal tissues and stained for GFP (green), Casp-3 (white), E-cad (magenta) and tdTom (yellow). Scale bar = 20 μm. D. FACS-purified GFP+ cells from s-SHIP-GFP/s-SHIP-Cre-tdTom mice were grown alone or in or co-culture with purified luminal cells, added at a 50:50 ratio and treated with 4-OH-Tam (5μM). After 7 days, they were fixed and stained for K5 and K8. Scale bar = 20 μm. E. Quantification of cell types in the cultures, measured as a percentage of tdTom+ lineage marked cells. n= 3 independent experiments. F. Schematic of the experiment. s-SHIP-GFP+ cells are cultivated on transwells with or without luminal cells at the bottom of the well. G. For co-culture experiments on Transwell plates, Luminal cells were added to the bottom of the well for 6-16 hours prior to adding GFP+ cells on the Transwell filter. Inserts were collected after 4 days. Scale bar = 50 μm. H. Quantification of cell types on the Transwell filters, measured as a percentage of tdTom+ lineage marked cells n=4 independent experiments (5,000 to 10,000 cells counted for each n). p-Values were determined by Sidak’s multiple comparison test (ns = not significant, *p< 0.05, **p< 0.01, ***p< 0.001, ****p< 0.0001).

### A short-range signal from luminal cells blocks cap cell multipotency

The in vivo luminal cell ablation experiments suggest that a repressive signal is generated by luminal cells that prevents cap cells from undergoing transdifferentiation to the luminal lineage. To test this hypothesis, we isolated GFP-tagged cap cells from our CreER;tdTom lineage tracing mouse crossed to the Tg11.5kb-GFP strain and incubated them in culture either alone or together with mature luminal cells. The tdTom marker was induced by addition of 4-OH Tamoxifen. After 1 week the cells were fixed, stained and tdTom+ cells were scored for K5 (basal marker) and K8 (luminal marker). About 60% of the GFP+ cap cells grown alone gave rise to K8+ or K8+/K5+ dual positive cells, consistent with our data from mammospheres (Fig 3D, E). Notably, however, in the co-culture almost 100% of the cells retained K5+ staining, and almost none expressed K8 (Fig 3D, E). We conclude, therefore, that mature luminal cells can signal to cap cells to suppress transdifferentiation, consistent with our in vivo ablation data (Fig 3C). Note that the added luminal cells form islands interspersed among the cap cells, so they are very close but not necessarily in direct contact with the cap cells (Fig 3D), suggesting that the luminal cells might release a soluble factor. We tested this possibility by using filter cultures in which GFP+ cap cells were grown on a filter above a layer of luminal cells, or in their absence (Fig 3F). However, in this situation the luminal cells had no effect on cap cell transdifferentiation (Fig 3G, H). This result suggests that the signal generated by the luminal cells is of short-range since it works on neighboring cap cells but not those separated by several millimeters of medium.

### An inhibitor screen reveals that TGFβ signaling regulates cap cell identity

To gain insight into the mechanism that determines whether cap cells function as myoepithelial progenitors versus transdifferentiation to the luminal lineage, we sorted pure GFP+ cap cells from Tg11.5kb-GFP transgenic mouse mammary glands and grew them in culture for 4 days in the presence or absence of signaling inhibitors (Table 1). After this time about 40% of the cells were still GFP+, while 40 - 60% had begun to express K8 (Fig 4A, B). Surprisingly, inhibitors against the WNT, Notch, JNK, NFkB, BMP, Hedgehog, mTOR, and YAP pathways had no significant effect on the proportion of GFP+ and K8+ cells. However, the STAT3 inhibitor Cucurbitacin I and JAK2 inhibitor Fedratinib, the PI3K inhibitor LY294002, and to a lesser extent the MEK inhibitor U0126, all promoted K8 expression and reduced GFP expression. Conversely, the ROCK inhibitor Y-27632 and TGFβR1 inhibitor Galunisertib potently inhibited expression of K8 and promoted retention of GFP expression (Table 1, Fig 4A, B). Galunisertib treatment also increased the number of cells in the culture (Supp Fig 5C). Because the initial cultures were 100% GFP+/K8-cells, these results strongly suggest that TGFβ and STAT signaling involve autocrine loops.

**Figure 4.**
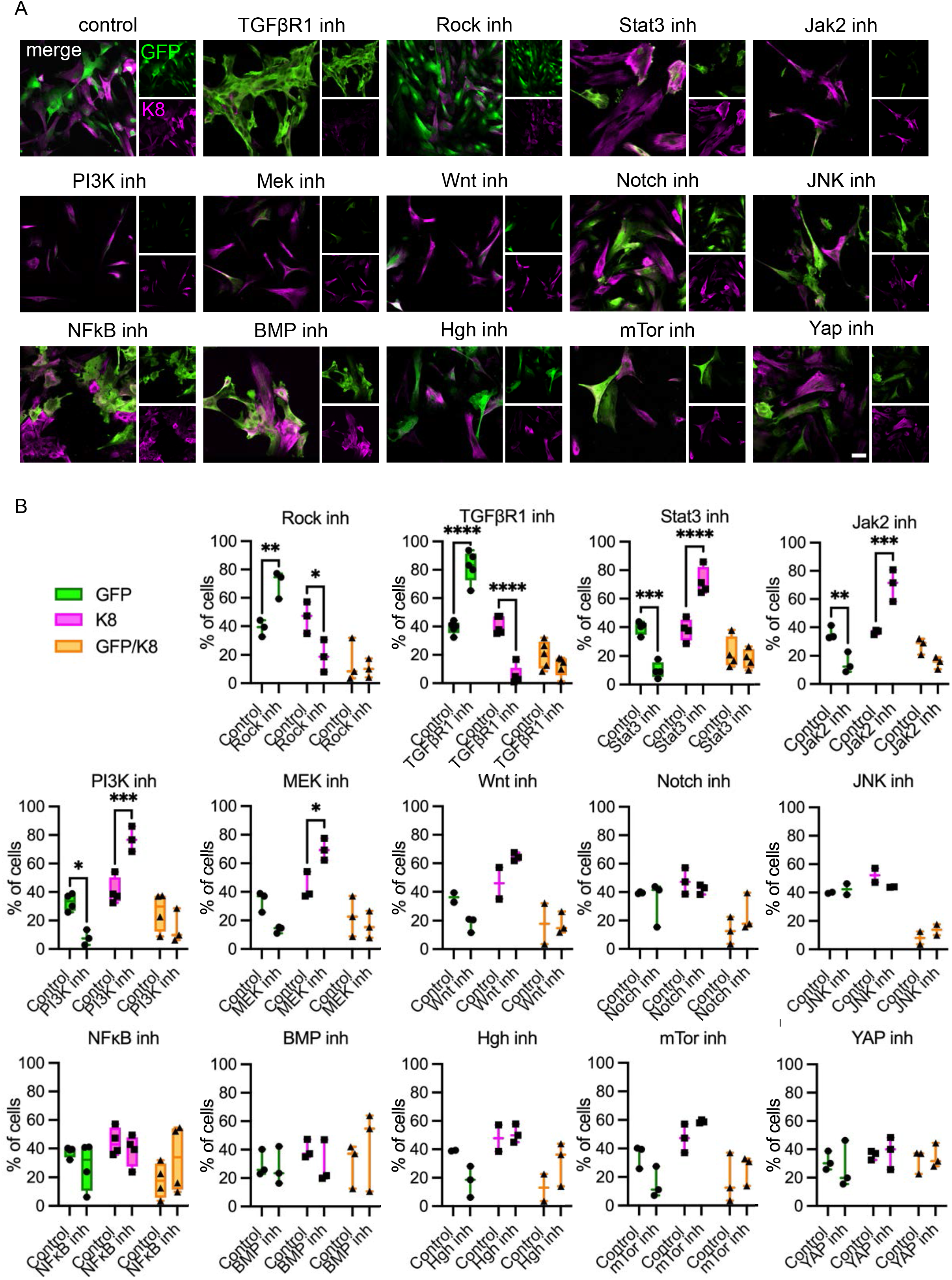
Screening for signaling pathways modulating cap cell fate. A. FACS-isolated GFP+ cells were plated on laminin and treated for 4 days with an array of different signaling pathway inhibitors, as listed in Table 1, then immunostained for GFP and K8. Scale bar = 50 μm. B. Percentage of cells respectively positive for GFP and K8, compared with the control. n=3 to 5 independent experiments (500 to 10,000 cells counted for each n). p-Values were determined by Sidak’s multiple comparison test (*p< 0.05, **p< 0.01, ***p< 0.001, ****p< 0.0001).

**Table 1.**

| Pathway | Inhibitor name | Concentration | Description | Effect on s-SHIP cells |
| --- | --- | --- | --- | --- |
| TGF-beta/<br>SMAD | Galunisertib<br>(LY2157299) | 10μM | TGFβ receptor 1 inhibitor | yes |
| TGF-beta/<br>SMAD | LY2109761 | 10 μM | Selective TGFβ receptor type<br>1 and 2 inhibitor | yes |
| RHO/ROCK | Y-27632 | 10μM | Inhibits ROCK1 and 2 | yes |
| JAK/STAT | Cucurbitacin 1<br>(JSI 124) | 100nM | Inhibits STAT3 and JAK2<br>activation | yes |
| JAK/STAT | Fedratinib<br>(SAR302503,<br>TG101348) | 1μM | Selective inhibitor of JAK2 | yes |
| PI3K | LY294002 | 25μM | Selective inhibitor of PI3K | yes |
| MEK | U0126 | 10μM | Selective inhibitor of MEK1<br>and MEK2 | yes |
| Wnt | LGK974 | 1μM | Porcupine inhibitor IV | no |
| Notch | DAPT | 2μM | γ-secretase inhibitor | no |
| JNK | JNK-IN-8 | 100nM | Inhibitor of JNK1,2 and 3 | no |
| NF-κB | JSH-23 | 10μM | Inhibits LPS-induced nuclear<br>translocation of p65 | no |
| BMP | LDN193189 | 1μM | Inhibits BMP type 1<br>receptors (ALK2 and 3) | no |
| Hedgehog | cyclopamine | 5μM | Inhibits Smoothened | no |
| mTOR | Rapamycin | 10nM | mTOR inhibitor | no |
| YAP | YAP-TEAD | 10μM | Potent and competitive<br>peptide inhibitor | no |
| Cyclin<br>Dependent<br>Kinase | Roscovitrine<br>(CYC202) | 10-40μg/ml | Potent and selective<br>inhibitor for Cdc2, CDK2 and<br>CDK5 | no |

To validate the effect of inhibiting the TGFβ pathway we used a second inhibitor, LY2109761, that targets both TGFβR1 and R2, which gave similar results with an increased fraction of GFP+ cells and strongly reduced numbers of K8+ cells (Fig 5A, B). We also used CRISPR/Cas-9 mediated gene editing to ablate the TGFβR1 and R2 receptors in isolated GFP+ cells, using a lentivirus, lentiCRISPRv2 to deliver the Cas9 and gRNAs. The cells were plated, selected for 24 hours with puromycin and assayed after 4 days. Efficiency of knockout was established by measuring the loss of mRNA with quantitative RT-PCR (Supp Fig 5A, B). Reduced levels of K8 expression and increased GFP were observed for deletion of each TGFβR gene, strongly supporting a role for TGFβ autocrine signaling in the cap to luminal cell conversion (Fig 5C, D). Addition of exogenous TGFβ1 reduced the number of GFP+ cells and raised the overall fraction of K8+ cells (Fig 5E, F). The small effect size is likely because the cap cells secrete TGFβ which activates receptors in an autocrine loop, so further raising the level of ligand has only a minor effect. Note that while receptor inhibition promoted cell proliferation the TGFβ-1 treatment reduced proliferation, raising the possibility that the observed changes in cell identity are linked to cell cycling (Supp Fig 5C, D). To test this hypothesis, we inhibited the cell cycle using Roscovitine, which significantly reduced cell number in primary cap cell cultures but had no effect on the percentage of cells undergoing luminal transdifferentiation (Supp Fig 5E-G). We conclude, therefore, that autocrine TGFβ signaling by cap cells drives their transdifferentiation independently of any effects on cell proliferation.

**Figure 5.**
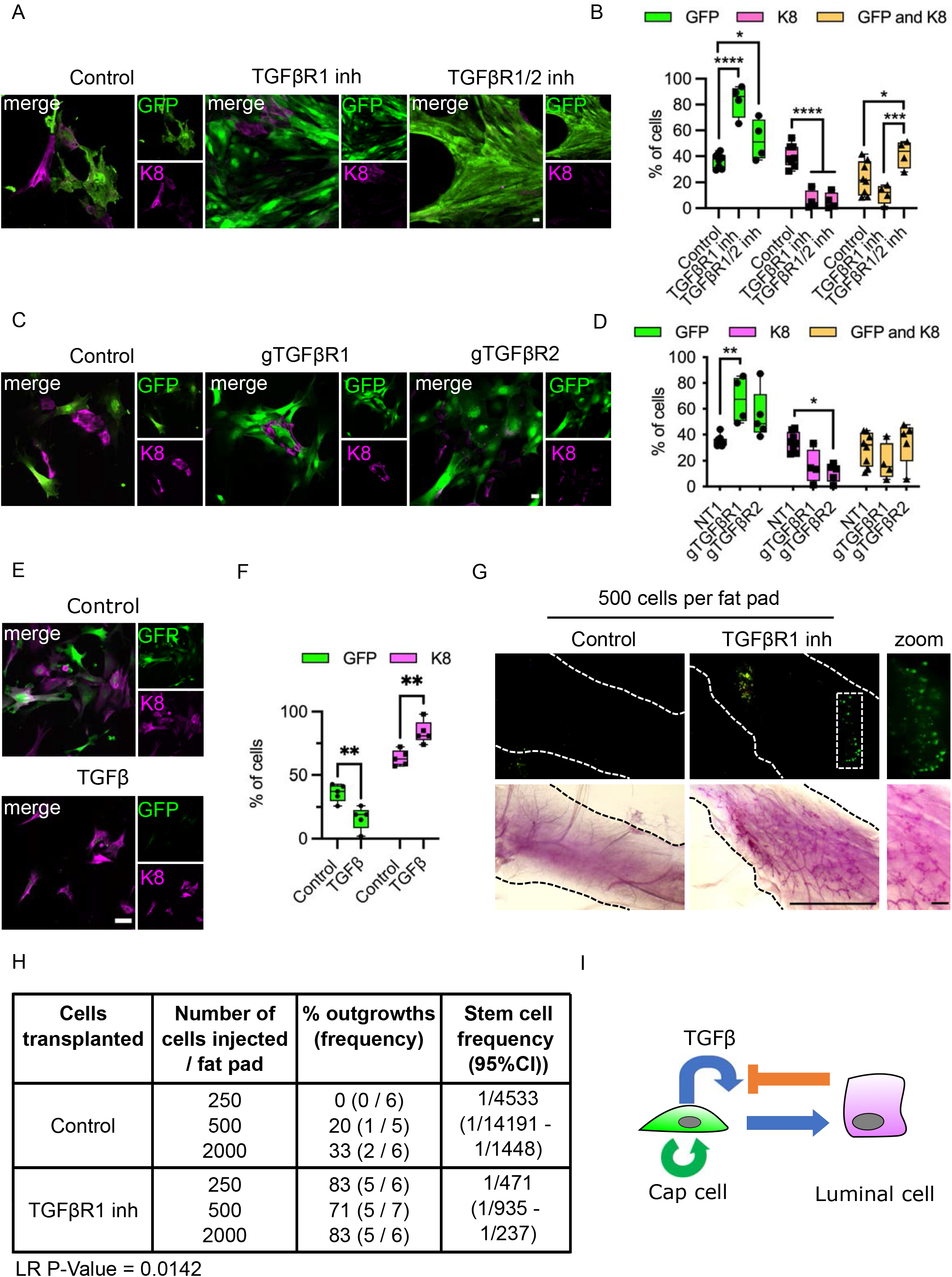
Blocking TGFβ Receptors maintains cap cell identity. A. FACS-purified GFP+ cap cells were plated on laminin, treated with TGFβ receptor inhibitors at 10 μM for 4 days, then fixed and stained for K8 and GFP. Scale bar = 50μm. B. Quantification of K8 and GFP positive cells compared with the control. n=5 independent experiments (1000 to 10,000 cells counted for each n). C. GFP+ cap cells were infected with lentiCRISPRv2 virus expressing guide RNAs gTGFβR1 or gTGFβR2 and selected 24 hours after with puromycin (0.5μg/ml). After 4 days they were stained as in A. Scale bar = 50μm. D. Quantification of K8 and GFP positive cells compared with the control. n=5 independent experiments. E. Effect of TGFβ on GFP+ cap cell identity. GFP+ cap cells were treated for 4 days with 10μg/ml recombinant mouse TGFβ1, then fixed and stained as in A. Scale bar = 50μm. F. Percentage of GFP+, K8+ and double positive cell ratio. n=4 to 5 independent experiments (500 to 10,000 cells counted for each n). p-values were determined by Sidak’s multiple comparison test (*p< 0.05, **p< 0.01, ***p< 0.001, ****p< 0.0001). G. Fluorescence image and Carmine alumn stained whole-mount images of mammary glands after 6 weeks where FACS-sorted GFP+ cells were plated on laminin, treated with TGFβR1inh, cultivated for 4-5 days then transplanted in cleared fat pads at different cells ratio. Scale bar 5mm and 2mm. H. Table summarizing 3 independent transplant experiments and the engrafted gland efficiency. Likelihood ratio (LR) P-Value = 0.0142. I. Model of a self-limiting cell circuit between pubertal mammary cap and luminal cells. Cap cells, which can self-renew (green arrow) release TGFβ, which in an autocrine loop promotes transdifferentiation of the cap cells to the luminal lineage (blue arrows). Neighboring luminal cells suppress this response (orange bar), maintaining the cap cell identity.

As a measure of biological functionality, GFP+ cap cells can efficiently regenerate mammary glands when transplanted into a recipient mouse from which the endogenous ductal tree has been removed. Therefore, to test if the increase in GFP+ cells produced by TGFβR1 inhibition is merely an isolated effect on the s-SHIP 11.5kb promoter or reflects a real increase in functional cap cells, we performed transplantation assays using cells grown in culture +/− inhibitor for 4 days, then injecting 250 – 2000 cells per fat pad. After 6 weeks the mammary glands were isolated for whole mount staining. Strikingly, TGFβR1 inhibitor-treated cells exhibited a ~10-fold increase in regenerative capacity relative to control cells (Fig 5G, H). We conclude that inhibition of an autocrine TGFβ loop in cap cells maintains their identity and regenerative capacity, while disruption of this circuit unlocks their potential to transdifferentiate (Fig 5I).

## DISCUSSION

The embryonic cells of murine mammary glands express a hybrid basal and luminal signature and are multipotent, generating both myoepithelial and luminal populations to form the rudimentary ductal tree (Spike et al., 2012; Wuidart *et al.*, 2018). However, by birth these two populations have become self-sustaining and multipotent stem cells are no longer present. Myoepithelial cells and three distinct subgroups of luminal cells are each maintained by separate progenitors throughout puberty (Spike *et al.*, 2012; Van Keymeulen et al., 2017; Wang et al., 2017). Importantly, however, DNA damage, or DTA-driven ablation of luminal cells, promotes a wounding response, with reversion to embryo-like multipotency in which myoepithelial cells transdifferentiate to the luminal lineage (Centonze *et al.*, 2020; Seldin and Macara, 2020).

During the rapid expansion of the ductal tree during puberty, TEBs arise at the tips of the mammary ducts, which contain distinct progenitor populations – luminal-like body cells and basal-like cap cells. Whether cap cells are multi- or unipotent, and whether they can transdifferentiate, has remained unclear (Ballard *et al.*, 2015; Prater *et al.*, 2014). Using a new lineage-tracing mouse, we now definitively show that cap cells in situ are unipotent progenitors of the myoepithelial lineage. However, removal from their normal tissue environment exposes a latent capacity for multipotency, while the addition of purified luminal cells back to cap cells in culture blocked multipotency, suggesting that luminal cells either sequester a transdifferentiation-promoting factor or release an inhibitory factor in a self-limiting cell circuit. Moreover, ablation of luminal cells in situ by DTA also triggers the cap cells to express stemness and generate new luminal cells by transdifferentiation.

A signaling inhibitor screen identified TGFβ and STAT signaling as likely regulators of this process. We probed the TGFβ mechanism and found that either inhibition of TGFβR1/2 or ablation of these receptors by CRISPR-Cas9 mediated gene editing prevented luminal transdifferentiation of purified cap cells, through a mechanism independent of the factor’s effects on cell proliferation. Moreover, cap cells treated with TGFβR inhibitor retained a higher capacity to regenerate mammary glands after transplantation into recipient mice. Together, these results are consistent with an autocrine mechanism in which cap cells release TGFβ that binds to receptors on the cap cell surface to promote transdifferentation to the luminal lineage.

Neighboring luminal cells limit this process so as to block cap cell plasticity and maintain homeostasis. This type of self-limiting cell circuit explains both the in situ and in vitro data. Moreover, it is consistent with earlier studies showing that conditional deletion of the TGFβR2 gene in mammary epithelium causes lobular-alveolar hyperplasia and TGFβ implants in the mammary gland suppress TEB formation and ductal outgrowth (Forrester et al., 2005; Silberstein and Daniel, 1987). It will be interesting to determine if TGFβ signaling plays any role in embryonic mammary gland development, and in the regulation self-limiting circuits in other circumstances.

## MATERIAL AND METHODS

### Mice

The Vanderbilt Division of Animal Care (DAC) ensures that all mice within the Vanderbilt facility are monitored daily for health status. DAC also ensures the overall welfare of the mice, and provides daily husbandry that includes environmental enrichment, clinical care, protocol record keeping, building operations, and security. The Vanderbilt mouse facility has three experienced Animal Care Technicians who attend to the daily needs of the animals. DAC ensures that all federal, state, and university guidelines for the care and use of animals are understood and maintained. Mice were housed with a standard 12 h light/12 h dark cycle. Mice were provided normal laboratory chow and water. Rosa-DTA (JAX stock #010527), Rosa-Tomato (JAX stock #007909) and K8.tGPD (JAX stock #032555) strains were obtained from the Jackson Laboratory (Bar Harbor, ME). s-SHIP-GFP mice were provided by L. Rohrschneider (Fred Hutchinson Cancer Research Center, Seattle, WA) (Bai and Rohrschneider, 2010). The s-SHIP 11.5kb promoter-CreER mouse was generated as described below and crossed to the tdTomato mouse to enable lineage tracing. All mouse experiments were performed with approval from the Vanderbilt Institutional Animal Care and Use Committee.

### Generation of s-SHIP-mERCremER mice

The Cre open reading frame was fused to 2 copies of a mutant Estrogen receptor (mER) ligand-binding domain (plasmid supplied by Chin Chiang, Vanderbilt University) (Zhang et al., 1996) and inserted downstream of the s-SHIP 11.5kb promoter (plasmid provided by L. Rohrschneider, University of Washington) (Rohrschneider et al., 2005). The DNA fragment containing the 11.5kb promoter, mERCremER and poly(A) signal was released from the backbone by Fsp1 digestion and microinjected into the pronuclei of B6 zygotes to generate 11.5kb-MerCreMer transgenic mice (Vanderbilt Genome Editing Resource) (Fig 1D). Three founders were identified by PCR, out of 31 mice born. Two were selected for subsequent studies.

### Induction of lineage tracing and cell ablation

Lineage tracing was induced at 4 weeks after birth for studies in puberty and pregnancy. A single pulse of tamoxifen (1.5mg; 50μl of tamoxifen (Sigma) at 30mg/ml in sunflower seed oil) was injected intraperitoneally. Tissues were collected after 2 days or 4 weeks for puberty studies and after one or three pregnancies. For DTA-mediated cell ablation experiments, 4-weeks old pubertal mice were induced with 9mg of tamoxifen (three injections of 3mg every 3 days). For s-Ship-CreER;K8-DTR mice, 6 mg of tamoxifen (two injections of 3mg 2 days apart) were injected at 4 weeks of age followed by a single DTA injection (5ng/g BW) at 5-weeks and tissue was collected at 6 weeks of age.

### Primers and gRNAs

All genotypes were performed by Transnetyx (Cordova TN). s-Ship-MerCreMer primer sequences were: Fw 5’-TTAATCCATATTGGCAGAACGAAAACG −3’; Rv 5’-CAGGCTAAGTGCCTTCTCTACA, s-Ship-GFP primer sequences were as described previously (Bai and Rohrschneider, 2010).

gRNAs were designed using CHOPCHOP. TGFβR1 and 2 primer sequences were as follows: gTGFβR1 Fw 5’-CACCGTTGACCTAATTCCTCGAGA −3’; gTGFβR1 Rv 5’-AAACGTCTCGAGGAATTAGGTCAAC −3’; gTGFβR2 Fw 5’-CACCGACCTGCAGGAGTACCTCACG −3’; gTGFβR2 Rv 5’-AAACCGTGAGGTACTCCTGCAGGTC −3’. The non-target gRNA sequence was as previously described (Fomicheva and Macara, 2020). gRNAs were inserted in the lentiCRISPR v2 plasmid (Addgene) (Sanjana et al., 2014). Lentiviral production was performed as previously described (McCaffrey and Macara, 2009) and viruses were concentrated using Lenti-X Concentrators (Takara).

### Antibodies, inhibitors and recombinant proteins

The following antibodies were used for immunostaining: anti-GFP(chicken, 1:1000, Abcam), anti-α-tubulin (mouse, 1:500, Sigma-Aldrich), anti-K14 (chicken, 1:500, Biolegend), anti-K5 (rabbit, 1:200, Abcam), anti-p63 (rabbit, 1:200, Cell Signaling), anti-E-cadherin (rat, 1:500, Thermofisher), anti-K8 (rat, 1:500, DSHB) and Hoechst. The following inhibitors were used: Jak2/Stat3 inhibitor (Cucurbitacin I, 100 nM, Tocris), Jak2 inhibitor (Fedratinib, 1 μM, Selleck Chemicals), TGFβR1 inhibitor (Galunisertib, 10 μM, Selleck Chemicals), TGFβR1/2 inhibitor (LY2109761, 10 μM, Selleck Chemicals), Rho/Rock inhibitor (Y-27632, 10 μM, Sigma-Aldrich), Hedgehog inhibitor (Cyclopamine, 5 μM, Tocris), Wnt inhibitor (LGK974, 1 μM, Cayman Chemical Company), Notch inhibitor (DAPT, 2 μM, Tocris), NF-κB inhibitor (JSH-23, 10 μM, Santa Cruz Biotechnology), BMP inhibitor (LDN193189, 1 μM, Tocris), JNK inhibitor (JNK-IN-8, 100 nM, Sigma-Adrich), mTor inhibitor (Rapamycin, 10 nM, Tocris), PI3K inhibitor (LY294002, 25 μM, Cell Signaling Technology), MEK inhibitor (U0126, 10 μM, Tocris), YAP-TEAD inhibitor (YAP-TEAD inhibitor 1, 10 μM, Selleck Chemicals), cyclin dependent kinase inhibitor (Roscovitine, 10-40μg/ml, Selleck Chemicals). Recombinant protein mouse TGFβ1 was from R&D systems and used at 10 ng/ml.

### RNA isolation and RT-qPCR

The following primers were used: GAPDH Fw 5’-GACCCCTTCATTGACCTCAAC-3’; Rv 5’-CTTCTCCATCGTGGTGAAGA-3’, TGFβR1 Fw 5’-CAGCTCCTCATCGTGTTGGTG-3’; Rv 5’-GCACATACAAATGGCCTGTCTC-3’, TGFβR2 Fw 5’-CCTCACGAGGCATGTCATCAG-3’; Rv 5’-ACAGGTCAAGTCGTTCTTCACTA-3’ (Oba et al., 2018).

RNA was isolated using RNeasy kit. cDNA was synthesized using SuperScript III Reverse Transcriptase, mixed with Maxima SYBR Green master mix. qPCR was performed on BioRad CFX96 Thermocycler. The relative gene expression was calculated from the average of technical triplicates, normalized to GAPDH and expressed using the ΔΔCt formula.

### Mammary cell isolation, cell staining and flow cytometry

The fourth and third mammary glands were dissected from 5.5 to 6.5-wk-old female mice, minced with scissors and digested in Digestion medium (DMEM/F12, 2mg/ml Collagenase A (Roche), 5μg/ml insulin (Sigma-Aldrich), 600 U/ml Nystatin (Sigma-Aldrich), 100 U/ml penicillin/streptomycin) for 2 hours at 37°C with periodic mixing. Epithelial organoids were collected by centrifugation at 1000rpm for 5 min. The cell pellet was resuspended into HF buffer (HBSS, 2% fetal bovine serum, 10mM Hepes pH 7.2) and NH_4_Cl (StemCell Technologies). The cell pellet was recovered after 5 min of centrifugation at 1000rpm and resuspended in organoid medium for organoid culture. For single cells, the cell pellet was dissociated in trypsin/EDTA (0.25%, Gibco), diluted in HF buffer, centrifuged at 1000rpm for 5 min. Pellet was resuspended in Dispase solution (StemCell Technologies) and DNase (1mg/mL, Sigma-Aldrich), dissociated into a single-cell suspension and filtered through a 40μm cell strainer (VWR). FACS analysis and cell sorting were performed using Fortessa and AriaIII instruments (Becton Dickinson). Antibody incubations were executed at 4°C for 10 min. Pacific Blue anti-mouse CD31 (rat, clone 390), anti-mouse CD45 (rat, clone 30-F11), anti-mouse Ter119 (rat, clone TER-119) were from BioLegend; APC anti-mouse CD326 or EpCAM (rat, clone G8.8) and PE-Cy7 anti-human/ mouse CD49f (rat, clone eBioGoH3) were from eBioscience. DAPI(1μg/ml) was added before analysis to exclude dead cells. The Lin^-^ population correspond to CD31, CD45 and Ter119.

### In vitro organoids and single cells assay

Single cells or organoids were cultured for one week in DMEM/F12 containing B-27 (Gibco), mouse EGF and mouse FGF basic protein (R&D Systems) and 50% of Matrigel (Corning). 4-OH-Tamoxifen (5μM) was added to the medium 24 hours after cells were plated in 2D culture. 4-OH-Tamoxifen (5μM) was added immediately in Matrigel with organoids or single cells. FACS-purified luminal and GFP+ cells were added at a 50:50 ratio, for co-culture experiments. For co-culture experiments on Transwell plates, luminal cells were added to the bottom of the well for 6-16 hours prior to adding GFP+ cells on the Transwell filter.

### Inhibitor assays

For inhibitor experiments, single cells isolated from 11.5kb-GFP mice were plated on laminin (20ug/mL, Sigma-Aldrich) and cultured in DMEM/F12 containing B-27 (Gibco), mouse EGF and mouse FGF basic protein (R&D Systems). Inhibitors (Table 1) were added 2 hours after cells were plated and fixed after 4 days in culture.

### Immunostaining and confocal microscopy

Dissected mammary glands were fixed 6 hours in 4% paraformaldehyde at room temperature. Tissues were washed in PBS and incubated overnight in 30% sucrose at 4°C. Tissues were embedded in OCT and kept at −80°C. Sections of 50μm were obtained using a Leica CM1950 Cryostat (Leica Biosystems). Tissue sections, cells or organoids were stained overnight, and images were acquired using Nikon A1R line scanning or Nikon spinning disk confocal microscopes, Nikon NIS Elements imaging software, a 20X/0.75 numerical aperture objective (NA), or 40X/1.30 NA objective with type B Immersion oil (Cargille Laboratories, Cedar Grove, NJ).

### Mammary transplants

FACS sorted-s-SHIP-GFP cells treated with or without TGFβR1 inhibitor for 4 days were dissociated in trypLE Select (Gibco) for 5 min. 2000-250 single cells in 10 ul of injection medium (20% Matrigel (Corning), DMEM/F12, 40 ng/mL EGF, 20 ng/mL (R&D Systems), FGF2 (R&D Systems), 0.01% trypan blue) was injected into the 4th cleared fat pads of 3-week-old C57Bl6 female mice (Jackson Laboratories) using a 26 needle gauge and Hamilton syringe.

### Mammary gland whole mounts

After 6-8 weeks of mammary ductal outgrowth, mammary fat pads were removed and fixed as described (McCaffrey and Macara, 2009). Pictures were acquired with an Olympus SZX16.

### Statistical analysis

All measurements were made in Fiji (Image J) or NIS-elements software. Cell angles of division were measured using Image J. Number of tdTom+ cells were calculated and classified as in the basal, luminal or cap cells compartment. Numbers of cells in cultures after treatment with Roscovitine, Galunisertib and TGFβ were counted using Hoechst as a cell marker, and normalized to the control. Numbers of GFP+ and K8+ cells were calculated using NIS elements software. All statistical analyses were performed using Student’s t-test, Sidak’s multiple comparison tes or two-way ANOVA test. s-SHIP-GFP stem cell frequency was calculated using a binomial generalized linear model with 95% confidence intervals and extreme limiting dilution analysis (ELDA) software (Hu and Smyth, 2009). The P-value was determined by the likehood ratio test.

## ACKNOWLEDGEMENTS

This work was supported by R35CA197571 from the National Cancer Institute, National Institutes of Health, DHHS (to IGM). Lineage tracing mice were generated in the Vanderbilt Genome Editing Ressources (VGER). The VGER is supported by the Cancer Center Support Grant (CA68485) and the Vanderbilt Diabetes Research and Training Center (DK020593). Flow Cytometry experiments were performed in the VMC Flow Cytometry Shared Resource. The VMC Flow Cytometry Shared Resource is supported by the Vanderbilt Ingram Cancer Center (P30 CA68485) and the Vanderbilt Digestive Disease Research Center (DK058404). Spinning disc confocal images were acquired using the Vanderbilt Cell Imaging Shared Resource (CISR).

## SUPPLEMENTAL FIGURE TITLES AND LEGENDS

**Figure S1.**
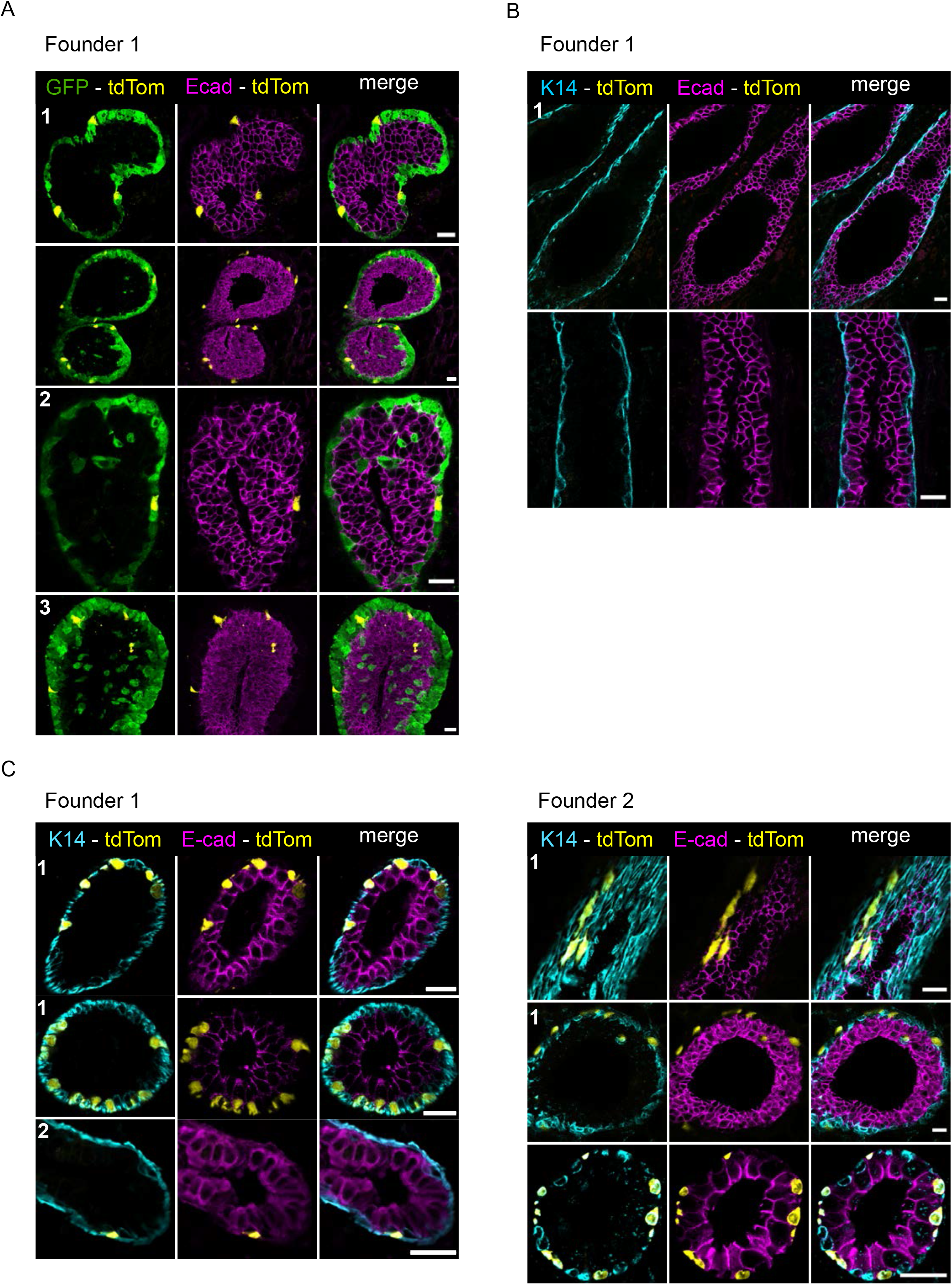
Specificity of the 11.5kb promoter CreER lineage tracing mouse for cap cells. A. and B. s-SHIP-Cre/tdTom mice were crossed with s-SHIP-GFP mice. Tamoxifen (1.5 mg/ml) was administered at 4-weeks-old and mammary glands were isolated after 2 days. Tissues were stained for GFP (green), K14 (cyan), tdTom (yellow) and, E-cad (magenta). Scale bar 20 μm. C. A single dose of Tamoxifen (1.5 mg/ml) was injected into 4 weeks-old s-SHIP-Cre/tdTom mice (founders 1 and 2), tissues were collected after 4 weeks and stained for K14 (cyan), tdTom (yellow) and, E-cad (magenta). Each number (1-3) represent an independent experiment. Scale bar 20 μm.

**Figure S2.**
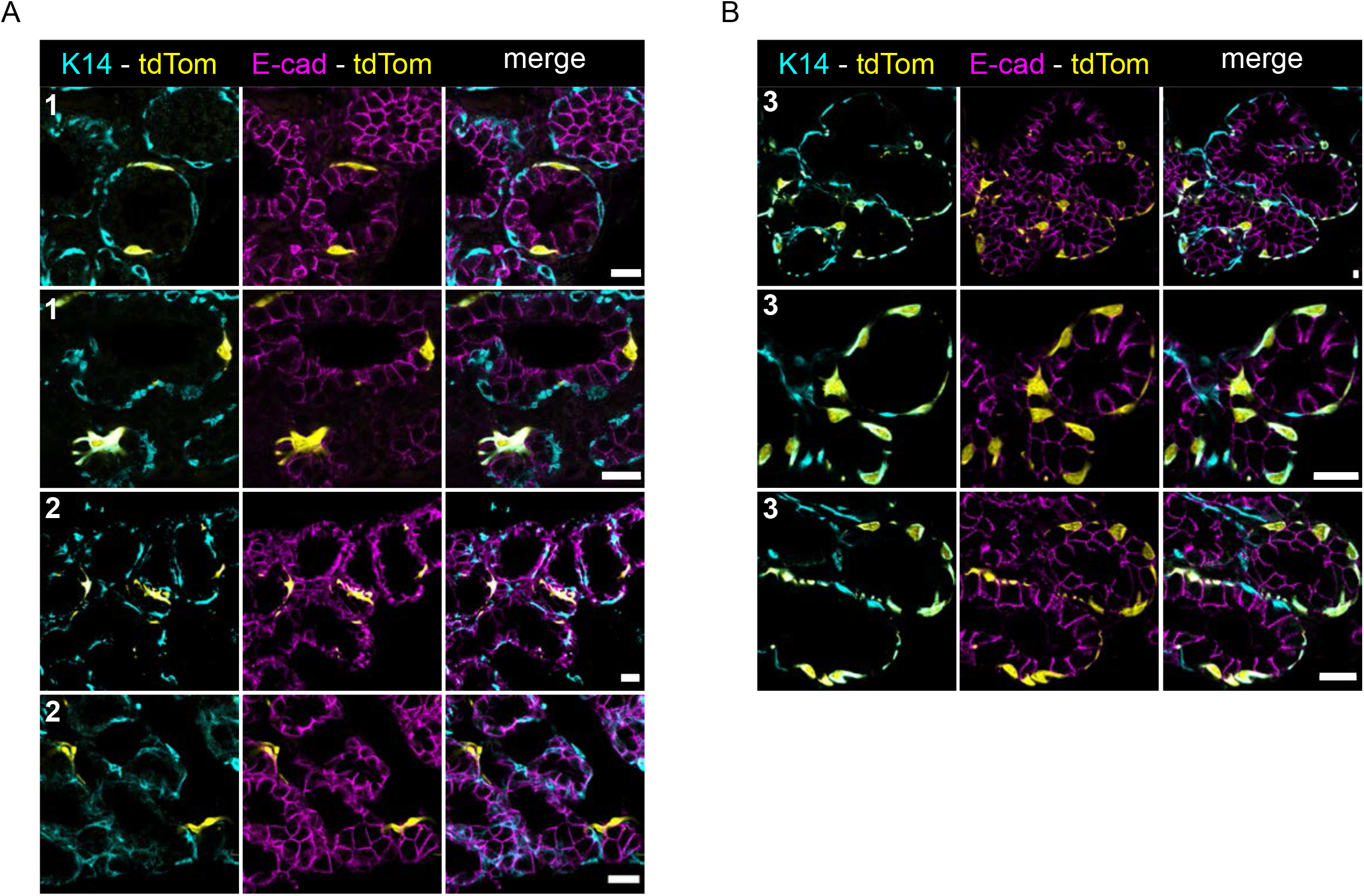
Alveoli from mice after 2 pregnancies show exclusively basal cell labeling with the tdTom lineage marker. A. and B. 4-weeks -old s-SHIP-Cre/tdTom mice were injected with 3mg/ml of Tamoxifen, mated at 7 weeks old and tissues were analyzed after one (A) or 3 (B) pregnancies. Each number (1-3) represent an independent experiment. Scale bar 20 μm.

**Figure S3.**
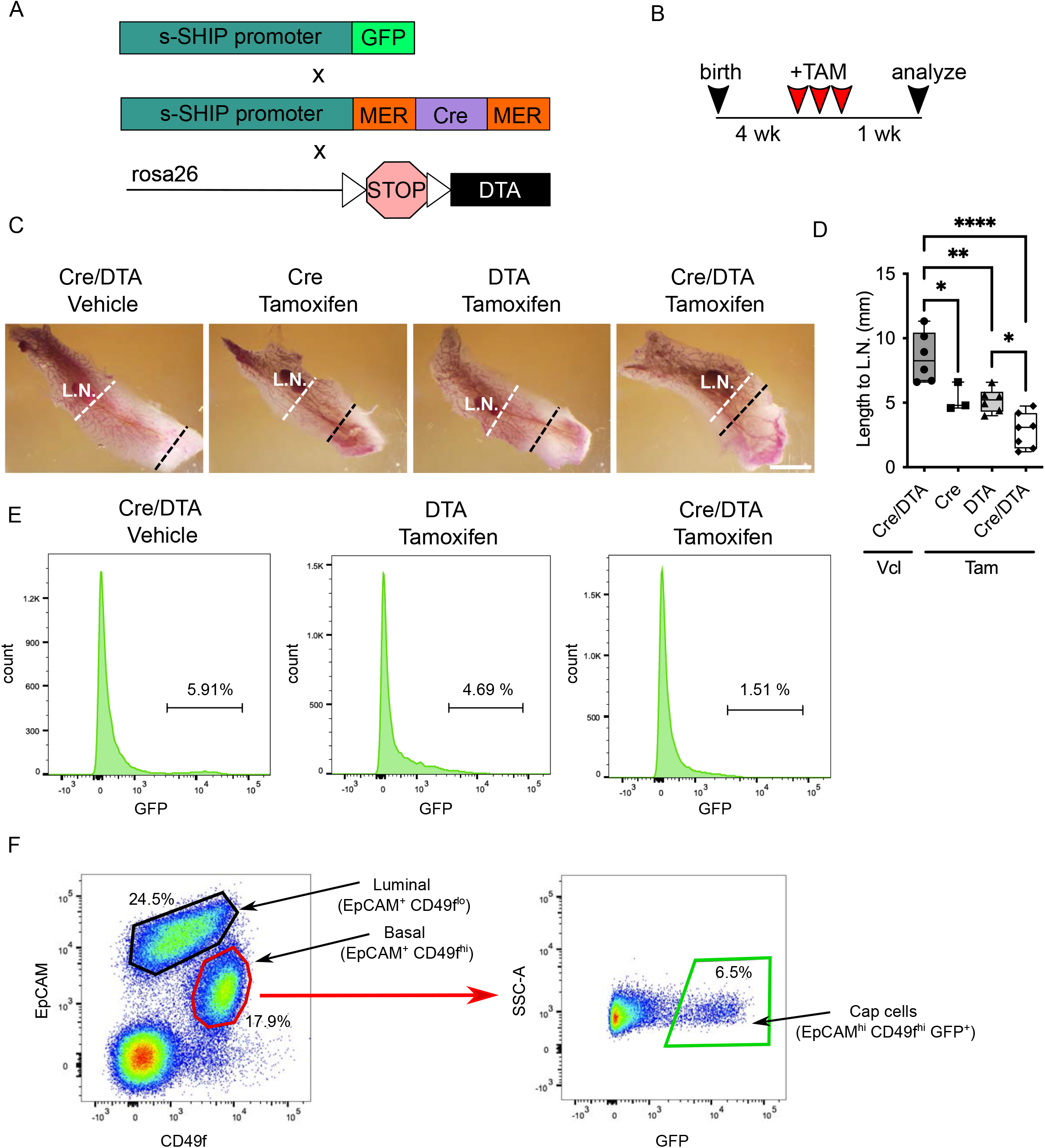
Cap cells are required for normal ductal outgrowth in the pubertal mammary gland. A. Schematic representation of the genetic strategy used to mark (GFP) and ablate cap cells. B. Protocol used to ablate cap cells with DTA. C. Image representation of Carmine alum-stained whole mount mammary glands. Scale bar 5 mm. D. Quantification of ductal tree length from lymph node (L.N.) to the tip of the leading edge of TEB. E. Cap cell numbers were quantified by FACS. A 4-fold decrease of GFP^+^ cells respective to the controls was observed after DTA-mediated ablation of the cap cells. F. GFP^+^ cap cells were isolated using standard gating strategy for flow cytometry analysis of mammary epithelial cells. Viable single cells and lineage negative for CD45/CD31/Ter119 were isolated. Representative FACS plot showing the luminal (EpCAM^+^ CD49f^lo^), basal (EpCAM^+^ CD49f^hi^) and cap (EpCAM^+^ CD49f^hi^ GFP^+^) subsets. The percentages represent the proportion of each cell subpopulation relative to their parent population.

**Figure S4.**
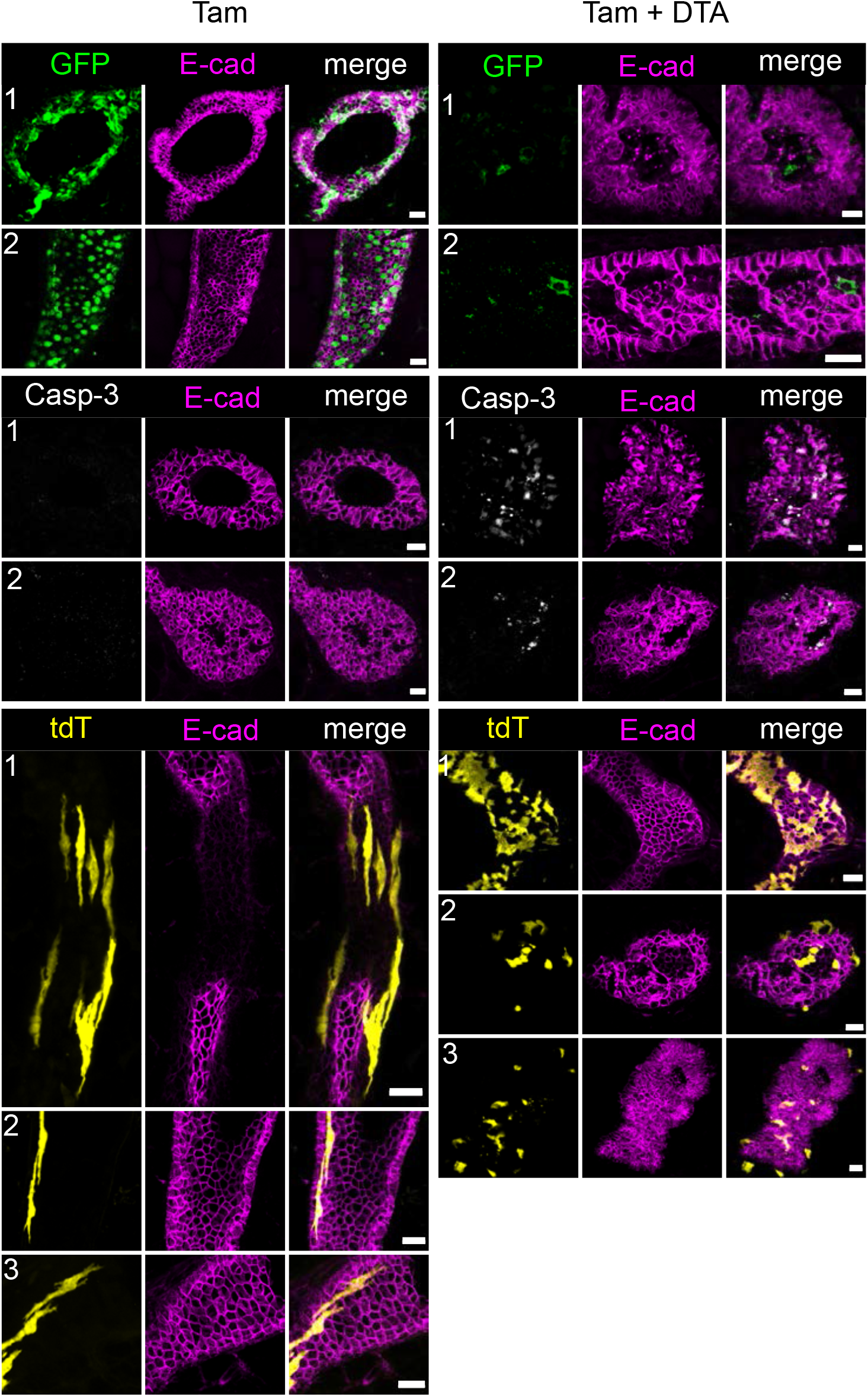
Ablation of luminal cells in situ triggers transdifferentiation of cap cells to the luminal lineage. s-SHIP-Cre/tdTom mice were crossed with K8-GFP-DTR mice. Mice were treated with 2 doses of Tamoxifen (3mg/ml) every 2 days then treated with a single dose of DTA (5ng/ml) one week after (Tam + DTA) or not (Tam). Tissues were collected one week after and stained for GFP (green), cleaved Casp-3 (white), tdTom (yellow) and E-cad (magenta). Each number (1-3) represent an independent experiment. Scale bar 20 μm.

**Figure S5.**
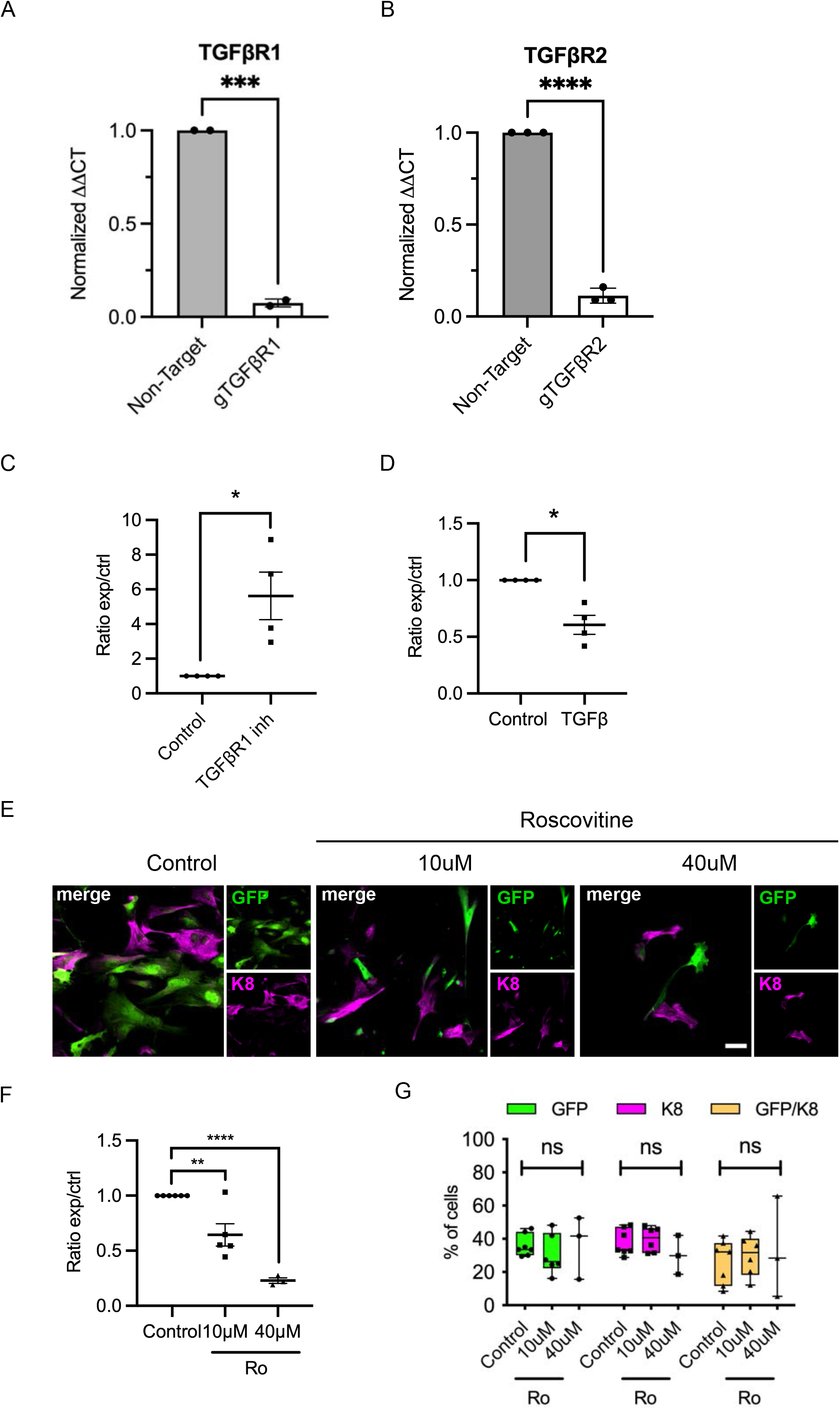
Expression of TGFβR mRNA after gene deletion, and of TGFβ ligands after siRNA-mediated knockdown, and lack of effect of cell cycle inhibition on cap cell identity. A, B. Relative gene expression levels of TGFβ receptors 1 and 2 obtained from control (Non-target) and TGFβ receptor 1 and 2 knockout cultures (nonclonal) of NMuMG cells. Samples were analyzed by qRT-PCR. ΔΔCt values from RT-PCR were normalized to the non-target control from 3 independent repeats. Error bar = Mean +/− S.D. Unpaired t-test was applied (*** p ≤ 0.001, **** p ≤ 0.0001). C, D. Ratios of cells treated with Galunisertib or recombinant TGFβ were quantified, and compared using a paired t-test (* p ≤ 0.05) E. Cap cells were sorted by FACS, cultured on laminin-coated plates and treated with roscovitine (Ro) at different doses (10μM and 40μM) for 4 days then stained for GFP and K8. Scale bar 50 μm. F. Numbers of cells were quantified by counting Hoechst-stained nuclei and normalizing to the control. Unpaired t test was applied (p ≤ 0.05, ** p ≤ 0.01, *** p ≤ 0.001, **** p ≤ 0.0001). G. Proportions of GFP+ and K8+ cells in cultures of GFP+ cap cells treated with Roscovitine. Percentages of cells positive for GFP and/or K8, compared with control (vehicle only). n=3 to 6 independent experiments and Turkey’s multiple comparisons test was applied (n.s. p> 0.05, * p ≤ 0.05, ** p ≤ 0.01, *** p ≤ 0.001, **** p ≤ 0.0001).

## Notes

### Competing Interest Statement

The authors have declared no competing interest.

